# Supramolecular fibrous hydrogel augmentation of uterosacral ligament suspension for treatment of pelvic organ prolapse

**DOI:** 10.1101/2023.01.08.522930

**Authors:** Beverly Miller, Wiley Wolfe, James L. Gentry, M. Gregory Grewal, Christopher B. Highley, Raffaella De Vita, Monique H. Vaughan, Steven R. Caliari

## Abstract

Uterosacral ligament suspension (USLS) is a common surgical treatment for pelvic organ prolapse (POP). However, the relatively high failure rate of up to 40% underscores a strong clinical need for complementary treatment strategies, such as biomaterial augmentation. Herein, we describe the first hydrogel biomaterial augmentation of USLS in a recently established rat model using an injectable fibrous hydrogel composite. Supramolecularly-assembled hyaluronic acid (HA) hydrogel nanofibers encapsulated in a matrix metalloproteinase (MMP)-degradable HA hydrogel create an injectable scaffold showing excellent biocompatibility and hemocompatibility. The hydrogel can be successfully delivered and localized to the suture sites of the USLS procedure, where it gradually degrades over 6 weeks. *In situ* mechanical testing 24 weeks post-operative in the multiparous USLS rat model shows the ultimate load (load at failure) to be 1.70 ± 0.36 N for the intact uterosacral ligament (USL), 0.89 ± 0.28 N for the USLS repair, and 1.37 ± 0.31 N for the USLS + hydrogel (USLS+H) repair (*n* = 8). These results indicate that the hydrogel composite significantly improves load required for tissue failure compared to the standard USLS, even after the hydrogel degrades, and that this hydrogel-based approach could potentially reduce the high failure rate associated with USLS procedures.

## 1. Introduction

Women’s health is an important, but chronically underfunded^[1]^ and therefore understudied, area of research. Increasing recognition of the unique aspects specific to women’s health^[2]^ has likely been influenced by the National Institutes of Health revised guidelines^[3]^ that required “women be included in all clinical studies” and “sex as a biological variable” to be considered beginning in 1994 and 2016, respectively. To address the substantial gaps in medical knowledge, researchers have leveraged tissue engineering approaches to transform our understanding of ovarian follicle development,^[4,5]^ endometriosis,^[6]^ gynecological cancers,^[7]^ and other disorders of the endometrium.^[8]^ In addition, special journal issues focusing on engineering for women’s health^[9–11]^ and in-depth review articles^[12,13]^ have increased visibility for this body of work. However, there is still much to be done to address the urgent clinical need to develop and investigate translational therapies.^[14]^ This is especially true for pelvic organ prolapse (POP) as surgical demand is predicted to increase by nearly 50% by 2050^[15]^ and outcomes from some treatment options, such as the uterosacral ligament suspension (USLS) procedure, remain discouraging.^[16]^

POP is a common condition affecting roughly 50% of parous women^[17]^ resulting in the descent of pelvic organs (vagina, bladder, uterus, small bowel, and rectum) due to compromised anatomical support structures.^[18]^ In healthy women, the levator ani muscles, the uterosacral ligaments (USLs), and cardinal ligaments support the pelvic floor organs via connective tissue attachments to the pelvis, while distal structures of the perineal body reinforce this supportive framework.^[19,20]^ Due to the relatively high failure rate of some native tissue repairs, an estimated one third of POP surgeries in 2010^[21]^ relied on non-absorbable lightweight polypropylene (PP) mesh. However, unacceptable post-surgical complications, such as mesh erosion, exposure, contracture and chronic pain, led the FDA to remove transvaginal mesh kits from the market in 2019.^[22]^ While PP mesh is still used by many pelvic surgeons to perform sacrocolpopexies, the current international climate and associated litigation surrounding the use of mesh leads many patients and their surgeons to seek mesh-free treatment options. The USLS procedure is a native tissue (suture only) alternative^[16]^ to mesh augmentation, but it is plagued by a failure rate of up to 40%.^[23,24]^ This procedure utilizes the dense collagenous sacral region of the USL to restore the vagina and surrounding structures to their anatomical position in the abdominal compartment via absorbable or non-absorbable sutures. According to recent studies,^[25]^ a woman’s current lifetime risk of undergoing prolapse surgery is approximately 20%,^[25]^ and that number is expected to rise in the coming years as the population ages. Therefore, there is an urgent clinical need to develop regenerative medicine strategies to provide safe and effective alternatives to mesh-based augmentation or suture only surgical procedures for POP.

There is growing interest in synthetic or biological materials to augment native tissue repairs.^[26,27]^ Namely, biodegradable polymers engineered to interact with the host tissue to promote constructive remodeling and integration^[28,29]^ could be a promising therapeutic platform for prolapse repair. Hydrogel biomaterials specifically provide an attractive alternative to mesh-based treatments due to the variety of material systems and chemistries available that can be tailored to the pelvic floor microenvironment. The ideal biomaterial for use in pelvic floor reconstructive surgeries is yet to be determined,^[30]^ but refinement of animal models can assist in the development of new materials. Several animal models including rats, mice, rabbits, sheep, swine, and non-human primates have been utilized^[31]^ in the study of pelvic organ prolapse with rodent models being optimal due to low cost and general accessibility. Despite this, the first POP surgical treatment rodent model was only recently established by our team.^[32]^ Previous studies investigating materials for prolapse repair in rats have used an abdominal hernia repair model^[33–38]^ due to the convenience of the established animal model^[39,40]^ and perceived conservation of outcomes between the abdominal wall and the pelvic floor.^[41,42]^ Conversely, while abdominal wall studies have demonstrated beneficial outcomes using increasingly stiff constructs,^[27,43]^ pelvic floor studies have found that increased stiffness is directly linked to the deterioration of vaginal smooth muscle and the surrounding pelvic floor.^[36,44]^ This “stress-shielding” phenomenon has also been seen in studies of bone^[45]^ as well as tendons and ligaments,^[46]^ where the stiffer material shields the adjacent tissue from experiencing physiological loads,^[47]^ causing the less stiff tissue to degenerate.^[30]^ Given this information, the abdominal wall hernia repair procedure is not the proven “proof of concept” model^[41,48]^ that it was once thought to be.

To address the need for alternative materials and a suitable animal model to investigate prolapse repair, our lab developed a hyaluronic acid (HA) based fibrous hydrogel^[49]^ as well as a rodent USLS model^[32]^ to investigate the augmentation of native tissue repair procedures. In this work, we present what we believe is the first study to use a rodent prolapse model to investigate a hydrogel biomaterial for the treatment of POP. The central aim of this study was to determine the impact of augmenting the USLS with a degradable hydrogel to restore mechanical integrity of the pelvic floor. A multiparous rat model was used due to its cost-effective nature and literature demonstrating the similar USL anatomy, cellularity, and matrix composition between rodents and humans^[50]^. Since failure of the native tissue repair is not well understood, the strength of the USL structures with their anatomical connections was assessed *in situ*. Our hypothesis was that the hydrogel augmentation of USLS would improve tissue integration at the USL-vaginal vault junction, therefore improving stability of the repair compared to sutures alone. Using a mechanical pull-off test,^[51]^ procedures that included the hydrogel augmentation demonstrated a significant increase in force required for failure compared to the USLS repair alone.

## 2. Results and Discussion

This study presents the experimental outcomes following investigation of a hydrogel biomaterial for augmentation of the uterosacral ligament suspension (USLS) procedure in rats. We utilized our lab’s recently developed rodent USLS model^[32]^ and supramolecularly-assembled injectable and photocurable fibrous hydrogel^[49]^ as a potential therapeutic for the treatment of POP. This is significant since there is an urgent clinical need for materials intended for the treatment of prolapse to be tested in pelvic floor models. While the mechanisms of mesh adverse events are not fully understood,^[44]^ it has become clear that the planes of fat, muscle, and fascia of the abdominal hernia repair rat model^[42]^ do not properly represent the smooth muscle and mucosa of the vagina and surrounding pelvic floor structures. Moreover, in addition to the USLS augmentation being more relevant for the investigation of potential prolapse therapeutics, previous work has demonstrated several similarities between rat and human USLs,^[50]^ further demonstrating the benefits of the rodent model. The rodent model also allows researchers to control several variables, such as parity, which we controlled for in this study. To the authors’ knowledge, this is the first study to investigate a potential non-mesh therapeutic for prolapse within the pelvic cavity.

### 2.1 Preparation of the injectable and fibrous hydrogel

Of the many classes of materials that have been used in tissue engineering, hydrogels have emerged as one of the most prominent and versatile as they can provide cells with a hydrated 3D environment that mimics native soft tissues.^[52]^ Polymeric hydrogels are especially attractive biomaterial platforms due to their tunable mechanical properties and ease of processing.^[53,54]^ With the stiffness of mesh-based augmentation implicated in vaginal tissue degeneration following implantation,^[44,55]^ there is an urgent clinical need for materials that can mimic mechanical properties of the pelvic floor tissues. We chose hyaluronic acid (HA) as our therapeutic biomaterial platform due to its hydrophilicity, degradability *in vivo*, and its role in supporting tissue repair.^[56]^ Here, the guest-host assembled hyaluronic acid (HA) hydrogel nanofibers were encapsulated in a matrix metalloproteinase (MMP)-degradable HA hydrogel that was delivered to the suture sites of the USLS procedure (**Error! Reference source not found**.). The guest (adamantane, Ad) and host (β-cyclodextrin, CD) modified fiber system leverages supramolecular chemistry that both enables facile injection for minimally invasive delivery and stabilizes the hydrogel post-injection. Both guest and host hydrogel fibers also included photocrosslinkable methacrylate groups for stabilizing the fibers following electrospinning. When the hydrogel fibers are mixed, the guest and host groups interact via hydrophobic non-covalent associations, creating a solid hydrogel until shear forces during injection disrupt the non-covalent interactions to create a “liquid-like” state. In Figure 1C, delivery of the hydrogel following hysterectomy is schematically shown where the hydrogel is applied over the sutures and surrounding tissue of the USL-vaginal vault created during the USLS procedure. For added stabilization of the self-assembling fibers within the abdominal cavity, the guest and host fibers were encapsulated in methacrylated peptide-modified HA (MePHA) engineered to be MMP-degradable, which allowed for secondary crosslinking via UV light.

**Fig. 1:**
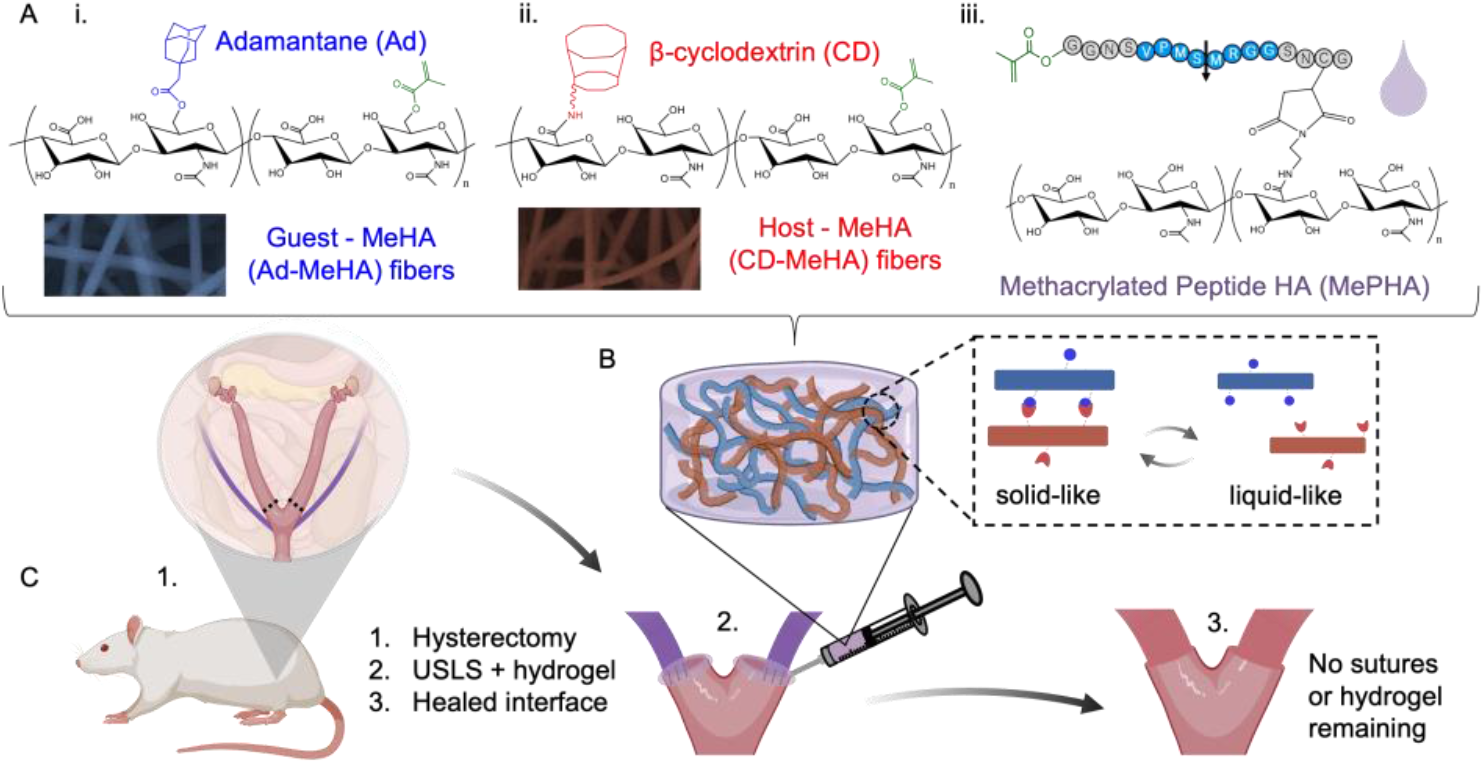
Schematic presentation of the composition of the fibrous injectable hydrogel system and uterosacral ligament suspension (USLS) augmentation process. A) Hyaluronic acid (HA) was modified with either i) the guest (adamantane, Ad) or ii) host (β-cyclodextrin, CD) molecules to create guest and host hydrogel fibers. Both guest- and host-modified HA also contained methacrylate groups to enable fiber stabilization via photocrosslinking after electrospinning. Separately, HA was functionalized with iii) a matrix metalloproteinase (MMP)-degradable photocrosslinkable peptide for encapsulating the fibers to produce B) a “solid-like” composite hydrogel capable of shear-thinning for injectable delivery to C) the sutures of the USLS procedure following hysterectomy. In situ photocrosslinking produced a mechanically stabilized biomaterial (shear modulus ∼ 4 kPa) that was fully degraded, as were the sutures, by the end of the study (24 weeks post-operative).

HA is an advantageous polymer choice in hydrogel design because it can be modified with reactive crosslinking groups to enable control of material biophysical properties (i.e., stiffness) and electrospun to create extracellular matrix (ECM)-like nanofibers mimicking native tissue.^[57–59]^ However, replicating host tissue properties following prolapse surgery is difficult without literature to define biomechanical properties. Studies done on the human prolapsed vagina report viscoelastic mechanical properties^[60]^ and an elastic modulus range of 2 – 13 MPa.^[61]^ The few studies that have investigated rodent pelvic support fixed the entire pelvic region and reported a linear stiffness of ∼ 3 N mm^-1^ for parous rats.^[62,63]^ To our knowledge, there have been no experimental studies conducted on the mechanics of the pelvic floor tissues following surgical treatment of POP, likely due to lack of available tissue samples and lack of prolapse repair animal models. Ideally, the hydrogel augmentation should enhance the properties of the structurally impaired vaginal tissue. Therefore, this study utilized our lab’s recently developed fibrous hydrogel platform^[49]^ as a first step in understanding ideal biomaterial design for the augmentation of prolapse repair. Oscillatory shear rheology (**Figure 2**) confirmed the hydrogel had robust mechanical properties under low strain conditions and underwent reverse gelation at higher strains mimicking injection prior to UV crosslinking. This is shown via a storage modulus that is higher than the loss modulus of 3.85 ± 0.61 kPa vs 0.65 ± 0.11 kPa respectively in conditions of low strain and a storage modulus that is lower than the loss modulus of 0.17 ± 0.01 kPa vs 0.53 ± 0.03 kPa respectively (*n* = 3). Importantly, the hydrogels showed full recovery of their viscoelastic properties after cyclic straining (Figure 2A). Once UV light was applied to the material, the storage modulus rapidly increased to 4.91 ± 0.51 kPa, demonstrating the stabilization of the hydrogel that would form over the suture tissue interface for the USLS augmentation (Figure 2B). Crosslinking was carried out with 365 nm light at an intensity of 10 mW cm^-2^ for 2 min.

**Fig. 2:**
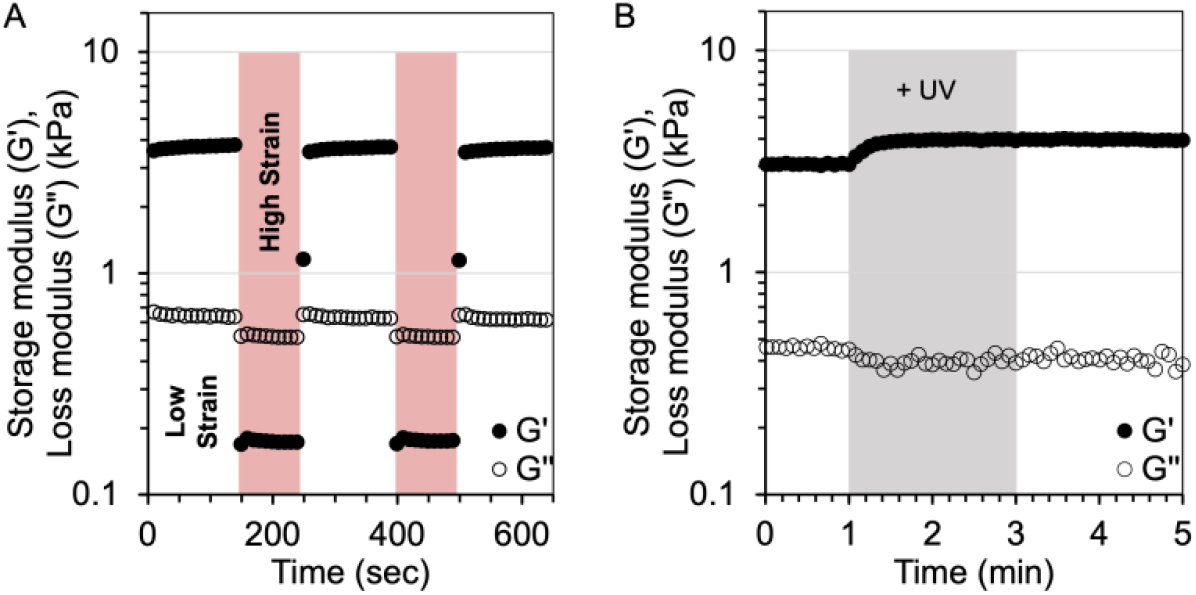
The mechanical properties of the fibrous hydrogel were measured using oscillatory shear rheology, showing A) a five-step strain sweep of low strain (0.5%, 100 s) and high strain (250%, 100 s) to illustrate the shear-thinning and self-healing properties of the hydrogel. Guest-host interactions between the fibers produced a solid-like material shown by the higher storage moduli than loss moduli at low strain. With the liquid-like behavior at high strain, shown by the storage moduli being lower than loss moduli, the material demonstrated shear-thinning and self-healing properties. B) In the presence of LAP photoinitiator, 2 min of UV light exposure (365 nm, 10 mW cm^-2^) increased the storage modulus of the material, indicating the crosslinking of the free methacrylates of the encapsulating MePHA hydrogel. All tests were performed at 37°C, *n* = 3.

### 2.2 *In vitro* biocompatibility

Many hydrogels formed from natural materials such as HA offer inherent biocompatibility.^[56,64]^ However, it is still important to demonstrate *in vitro* biocompatibility of any developed material system prior to use *in vivo*. Therefore, cytotoxicity and hemocompatibility assays were performed to evaluate the biocompatibility of the fibrous hydrogel composite. **Figure 3** shows the cytotoxicity of each hydrogel component (Ad-MeHA, CD-MeHA, and MePHA) when exposed to human mesenchymal stromal cells (hMSCs) in culture. There were no statistically significant differences for cell metabolic activity in the polymer groups compared with the control group incubated in media with PBS for 1 and 3 days. Metabolic activity, normalized to PBS control levels after one day, were shown to be 132.3 ± 25.1% for Ad-MeHA, 190.8 ± 33.5% for CD-MeHA, and 158.4 ± 23.1% for MePHA after one day of culture. Metabolic activity after three days increased to 179.9 ± 34.1% for Ad-MeHA, 193.6 ± 16.1% for CD-MeHA, and 185.8 ± 37.7% for MePHA (*n* = 9). These cytocompatibility results are in alignment with previous results using the fibrous hydrogel system, where hMSC viability was over 85% following injection and after 7 days of culture.^[49]^

**Fig. 3:**
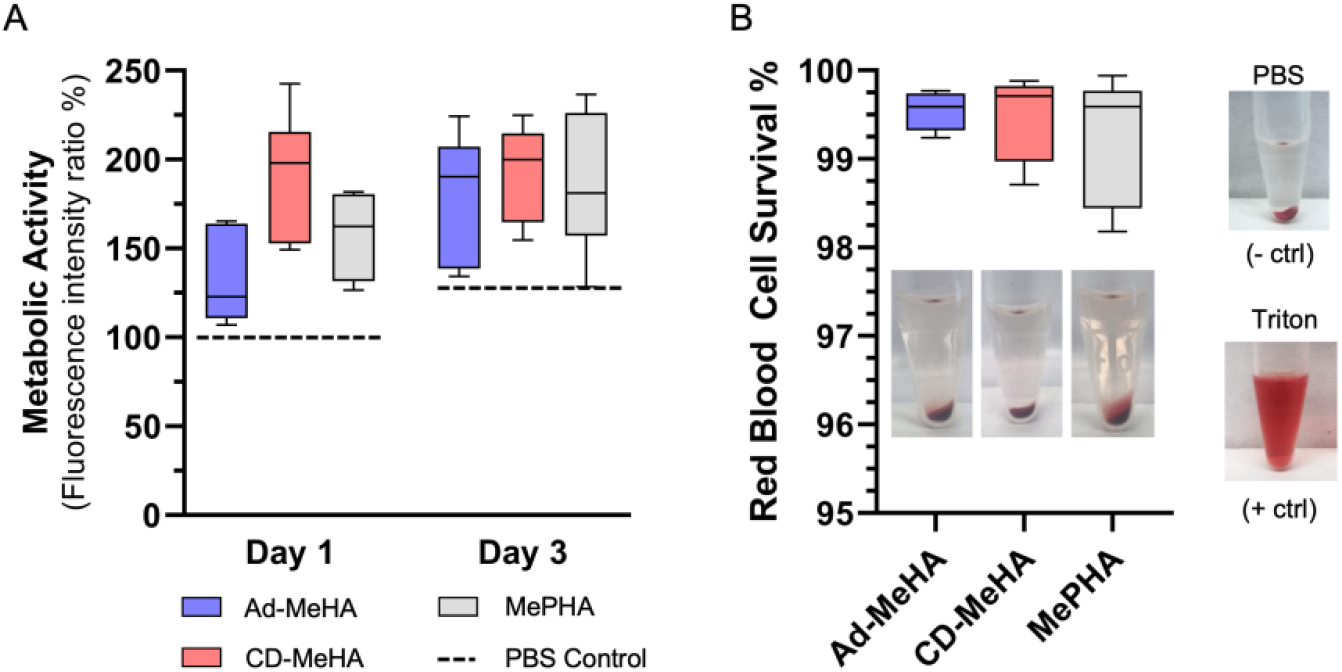
*In vitro* cytocompatibility and hemocompatibility tests. A) Metabolic activity, measured via an Alamar blue assay, of human mesenchymal stromal cells cultured in the presence of hydrogel components for 1 and 3 days compared to cells cultured with PBS (*dashed lines*). B) Percentage of rat red blood cell (RBC) survival following incubation with hydrogel components. Pictures of the RBCs after centrifugation include PBS (negative control) and Triton-X100 (positive control).

Unlike the *in vitro* studies, materials implanted *in vivo* will encounter the surrounding tissue as well as bodily fluids such as blood. While the fibrous hydrogel is designed to mimic the native ECM and support tissue function, interactions between the biomaterial and blood should also be evaluated prior to implantation. Therefore, assays that investigate the reaction of red blood cells (RBCs) with the scaffold provide useful information regarding possible consequences of biomaterial-blood interactions.^[65]^ Figure 3B shows the hemocompatibility for each component using freshly collected rodent RBCs. After incubation with the material components, the RBCs showed no obvious hemolysis compared to the Triton-X100 group (positive control). RBC survival rates were shown to be 99.6 ± 0.2 % for Ad-MeHA, 99.5 ± 0.4 % for CD-MeHA, and 99.3 ± 0.7 % for MePHA, with the overall material showing a survival rate of 99.4 ± 0.4 % (*n* = 9). The low hemolysis ratio indicates that the fibrous hydrogel scaffold possesses satisfactory blood compatibility. Together, these *in vitro* experiments demonstrated biocompatibility of the fibrous hydrogel composite.

### 2.3 Augmentation of rodent USLS

The overall body of tissue engineering literature for prolapse repair is dominated by mesh-based augmentation, with the implication that native tissue and mesh-based repairs are of the few suitable treatment options for POP. With this narrow focus on mesh-based therapeutics, it is unsurprising that the search for ideal tissue remodeling materials for urogynecological repair is ongoing. A lack of accessible, and surgically accurate, animal models is at least partially to blame for the lack of non-mesh biomaterial investigation. In fact, just last year Bickhaus and colleagues were the first to use a rodent model to investigate a new polycarbonate urethane mesh for pelvic reconstruction via implantation in the vagina.^[66]^ However, this study still focused on a mesh construct and used a surgical method not used in humans when repairing prolapse. By contrast, our rodent USLS procedure mimics the surgical method used in humans, establishing this work as the first study to investigate biomaterial augmentation for prolapse repair in a surgically accurate rodent model.

In this study, we aimed to investigate our fibrous hydrogel composite using our established rat USLS surgery model.^[32]^ All animals were maintained and treated under the approval of the University of Virginia Institutional Animal Care and Use Committee. All rats were multiparous, meaning that they had delivered prior litters (two in our study). Twenty-four Lewis rats (Charles River Laboratories) between 4 and 6 months of age were used to accommodate the two-litter requirement. Detailed surgical instructions with reproducible steps are published elsewhere.^[32]^ Briefly, following anesthetization using isoflurane and aseptic preparation of the surgical site, a vertical midline skin incision was made down the linea alba and the muscle layer underneath. Removal of the uterine horns (hysterectomy) created the vaginal vault structure with the intact USLs directly below (**Figure 4A**). Ovaries were left *in situ*. The USL-vaginal vault interface was created by suturing the remaining vaginal vault tissue to the USLs (Figure 4B). Animals were randomly assigned to one of the following experimental groups: sham surgery with no prolapse repair (USL; *n* = 8), prolapse repair (USLS; *n* = 8), or prolapse repair with hydrogel augmentation (USLS+H; *n* = 8).

**Fig. 4:**
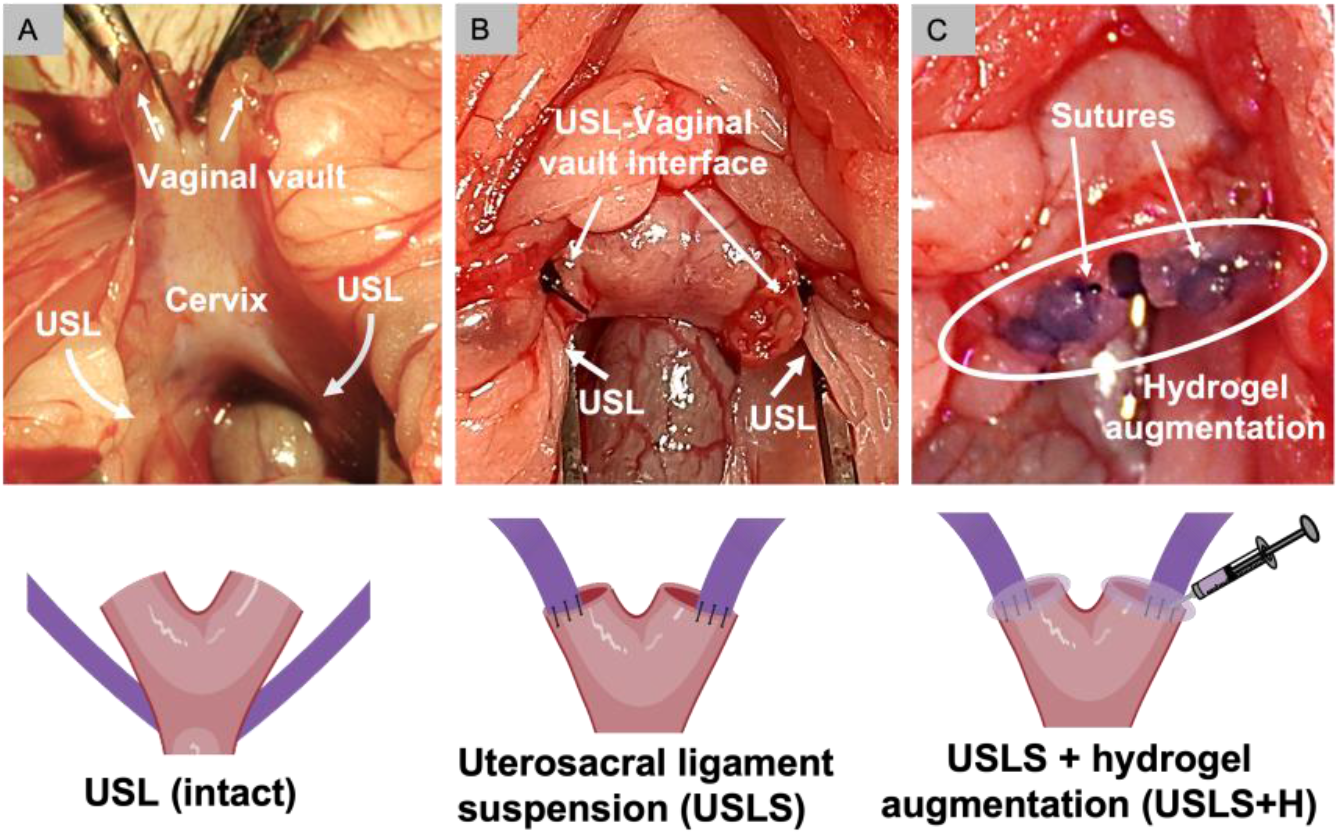
Schematic presentation of the uterosacral ligament suspension (USLS) surgical process and hydrogel augmentation (USLS+H). A) Removal of the uterine horns creates the vaginal vault structures with the USLs seen directly adjacent to the cervix. B) The USLS procedure secures the vaginal vault high on the USLs via a single suture such that the vaginal vault is elevated toward the sacrum. C) For hydrogel augmentation, 20 μL of hydrogel per side was administered such that the suture at the USL-vaginal vault interface was completely covered prior to 2 min of UV crosslinking to further stabilize the hydrogel.

For the animals randomly assigned to the USLS+H experimental group, the injectable fibrous hydrogel was administered and then UV crosslinked to stabilize the material over the suture (Figure 4C). Prior to delivery, the fibrous hydrogel composite was prepared using sterile technique. Each component (Ad-MeHA fibers, CD-MeHA fibers, MePHA) was sterilized by overnight lyophilization followed by germicidal UV irradiation for 2-3 hr. Additionally, the LAP photoinitiator stock solution was sterile filtered before use. The supramolecular nature of the guest-host fibers kept the 20 μL of hydrogel delivered to each suture site, total of 40 μL per animal, localized while the UV crosslinking further stabilized the material. We chose UV photopolymerization for *in situ* stabilization as it is a commonly used technique for covalent crosslinking of hydrogels.^[67–69]^ However, to utilize the minimally invasive capabilities of the fibrous hydrogel in another animal model or in human treatment, alternative *in situ* stabilization mechanisms such as redox-initiated polymerization would potentially need to be explored. Delivery of the hydrogel was not minimally invasive in this study, but the benefit of the self-healing property of the material cannot be overstated. The syringe and needle delivery allowed for precise placement and minimized migration of the material prior to UV stabilization. Material migration is a known issue when implanting biomaterials *in vivo*.^[52]^ Given the exposure of the implanted material to the abdominal cavity, the USLS model is no different. With these concepts in mind, other self-healing biomaterials may be good candidates to investigate for prolapse repair due to the benefits of localized and minimally invasive delivery.

### 2.4 *In vivo* assessment of hydrogel degradation

In developing materials for pelvic floor applications, biodegradability has been identified as a key consideration.^[26,36]^ *In vivo* erosion of the fibrous hydrogel scaffold was investigated by covalently attaching a near-infrared (NIR) dye to the fibers and monitoring the corresponding loss in signal intensity over a 6 week period (**Figure 5**). In addition, the results from the fluorescent imaging confirmed sustained placement of the hydrogel at the suture sites. A maleimide-modified cyanine 7.5 fluorophore (Cy7.5, Lumiprobe Corporation, Figure S1) was conjugated to a methacrylated peptide (Figure S2) to enable UV light-mediated methacrylate crosslinks to covalently attach the Cy7.5-labeled methacrylated peptide (Cy7.5-Me, Figure S3) to the hydrogel material. In Figure 5B the labeled hydrogel covers the sutures of the USLS procedure with Figure 5C showing serial *in vivo* optical images. The fibrous injectable hydrogel scaffold demonstrated faster erosion over the first week (losing more than 50% of the NIR signal) and ∼ 95% degradation after six weeks, showing an exponential decay erosion trend. The rapid degradation is hypothesized to be due to the small amount of material, the exposure to abdominal movements (physical) and fluid erosion (convective fluid movement), and cell-mediated enzymatic degradation. It is unclear if this degradation rate is ideal as there is a limited understanding of the remodeling rate at the USL-vaginal vault junction. General tendon and ligamental remodeling has a known timescale of a few weeks,^[70]^ but that does not necessarily represent the supportive ligaments of the pelvic floor. Meanwhile, there is no set time-scale reported for vaginal and pelvic smooth muscle remodeling, but there are clear examples of hormone^[71,72]^ and strain dependence.^[73]^ While only a semi-quantitative assessment of degradation, these results demonstrate the degradability of the fibrous HA composite scaffold and provide context for future studies investigating biomaterial degradation in POP models.

**Fig. 5:**
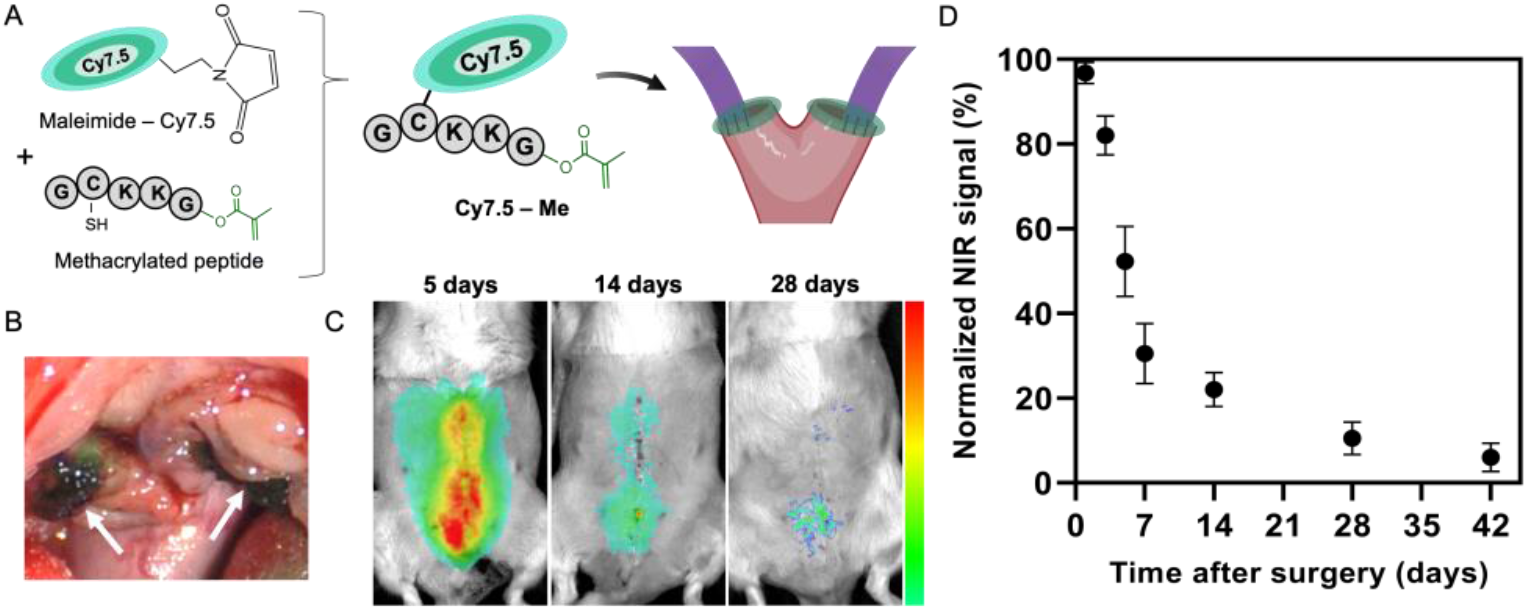
*In vivo* fibrous hydrogel degradation. A) Methacrylated peptide conjugated with sulfomaleimide cyanine 7.5 (Cy7.5-Me). The methacrylated peptide was first synthesized via solid-phase synthesis and then the Cy7.5 was tethered via maleimide-thiol click chemistry. The methacrylate on the peptide allowed for covalent conjugation of the Cy7.5 dye to the HA backbone via photocuring before scaffold formation. B) Arrows point toward the Cy7.5-labeled hydrogel following delivery. Erosion of the fibrous hydrogel, as represented by the C) serial *in vivo* optical images, was D) quantified by Cy7.5 signal decay (*n* = 4).

### 2.5 *In situ* assessment of specimen mechanical integrity 24 weeks post-operative

While urogynecological research is still in its infancy compared to other fields, this work builds on previous work^[61,74]^ attempting to provide biomechanical context for pelvic floor tissues. At 24 weeks post-operative, the mechanical properties of the USL-vaginal vault interface after suture only (USLS) and hydrogel augmentation (USLS+H) procedures were assessed and compared with mechanical properties of the intact the USL (**Figure 6**). Mechanical “pull-off” tests were performed *in situ* (Instron, 10 N load cell, 0.00025 N resolution). Specimens were preloaded at 0.15 N and then preconditioned at an elongation rate of 0.1 mm sec^-1^ as described in our previous work.^[32]^ Data points were collected every 0.1 sec and analyzed via MatLab. The sutures or the sutures + hydrogel were completely resorbed at the time of testing. Briefly, the USL-vaginal vault interface of the USLS and the USLS with hydrogel (USLS+H) groups were exposed such that umbilical tape could be placed beneath the structure and then clamped in the tensile tester grip. For the intact tissue comparison, the umbilical tape was placed beneath the intact USL structure. A total of 8 specimens (*n* = 8) are reported per group. After specimen preparation, samples were tested until failure where Figure 6B shows a sample mid-test and Figure 6C is an overview of defined parameters of the testing.

**Fig. 6:**
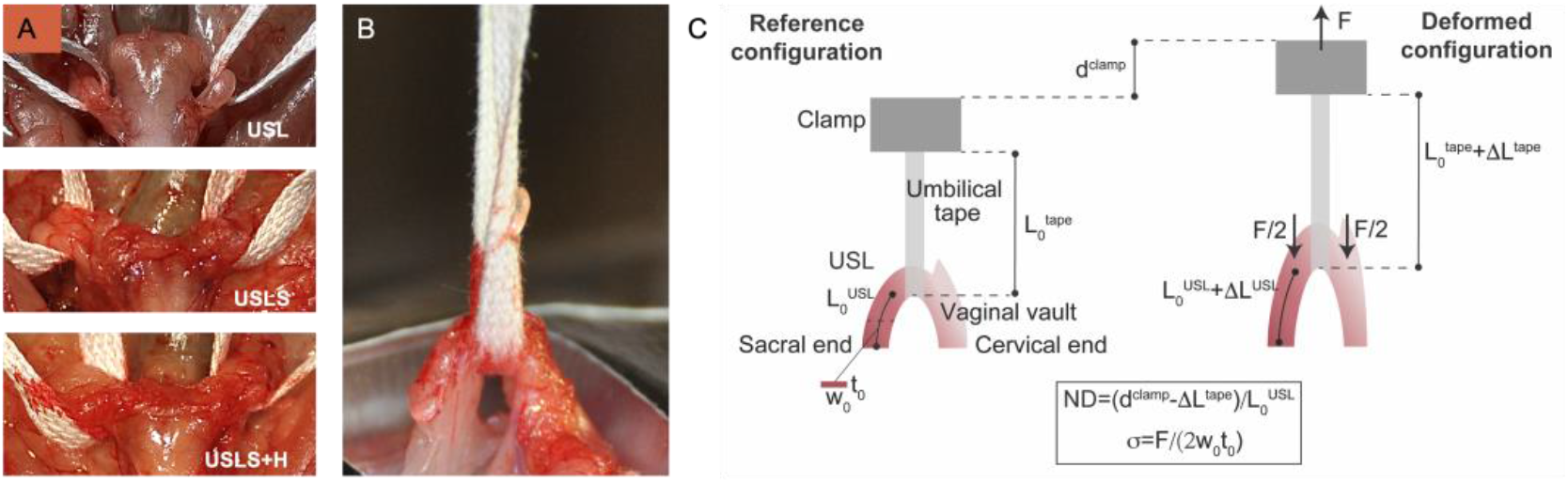
*In situ* tensile tests pulled to failure. A) Tissue preparation for tensile testing where the umbilical tape was placed underneath the cervical end of the structure. B) Sample undergoing the tensile “pull-out” test to failure with a schematic detailing the testing measurements during C) the reference configuration and the deformed configuration.

Load-displacement curves are reported in Figure S4 where displacement is defined as *d*^clamp^ – ΔL^tape^, where *d*^clamp^ is the displacement of the clamp (or cross-head displacement) and ΔL^tape^ is the change in length of the umbilical tape. The presence of several peaks in the load as the displacement increased demonstrated that the failure of the USLs, or the surgically repaired tissue, occurred gradually. As expected for soft tissues, data were characterized by an initial toe region, a more linear region, and then a nonlinear failure region where the gradual failure is likely due to bundles of tissue fibers breaking at various load levels.

Results showed an ultimate load of 1.70 ± 0.36 N for the intact uterosacral ligament (USL), 0.89 ± 0.28 N for the USLS repair, and 1.37 ± 0.31 N for the USLS+H group (**Figure 7**). The stiffness, defined as the steepest positive slope measured over a 1 mm elongation interval, was 0.48 ± 0.10 N/mm for the intact USL, 0.28 ± 0.07 N/mm for the USLS procedure, and 0.42 ± 0.10 N/mm for the USLS+H group. Stress data were computed under the assumption that the USL was loaded along one axis as shown in Figure 6C. Next, the ultimate stresses were calculated using the ultimate load data. Tissue measurements including the thickness (t_0_), width (w_0_), and gauge length (L_0_^USL^), where the gauge length was defined as the length of the USL from the sacral attachment to the vaginal vault/cervical insertion, were also obtained (Figure 6C). Average cross-sectional thickness and width were 1.9 ± 0.3 mm and 2.8 ± 0.2 mm while average gauge length was 12.3 ± 0.7 mm respectively. The ultimate stress in the tissue before complete failure was shown to be 0.30 ± 0.06 MPa for the intact USL, 0.17 ± 0.04 MPa for the USLS, and 0.27 ± 0.02 MPa for the USLS+H groups. Notably, for all mechanical testing data shown in Figure 7, hydrogel augmentation (USLS+H) resulted in statistically significant increases in mechanical properties compared to the USLS-only group. Additionally, there were no statistically significant differences in mechanical properties between the USLS+H and intact USL groups. These exciting results, obtained after the hydrogel and sutures were completely degraded, highlight the potential of this hydrogel therapeutic to promote long-term improvement in the mechanical stability of the USL-vaginal vault interface created in USLS procedures.

**Fig. 7:**
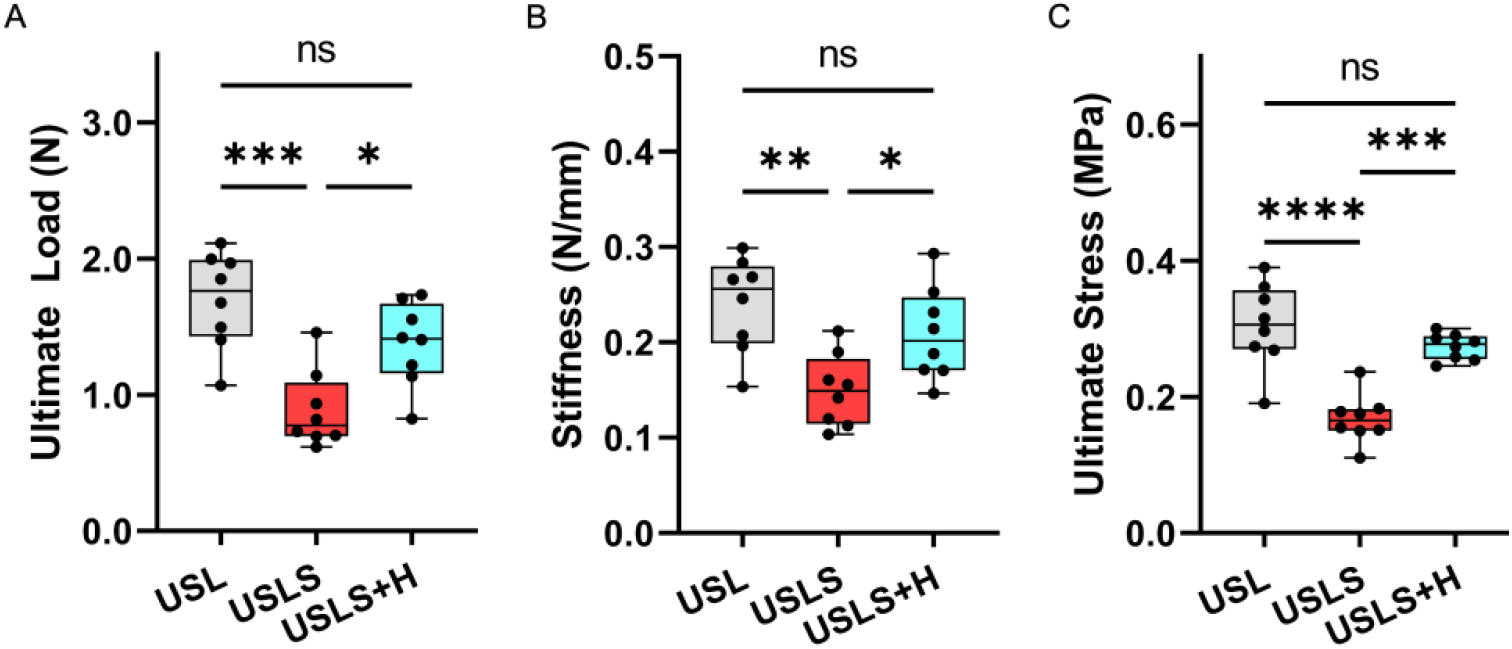
Analysis of sample mechanical properties based on the load-displacement data and calculated *stress*. Data presented are A) force at failure, or ultimate load, B) slopes of the load-displacement curves (stiffness), C) and the stress at failure, or ultimate stress. *n* = 8, * *P* < 0.05, ** *P* < 0.01, *** *P* < 0.001, **** *P* < 0.0001.

The mechanical testing protocol described here represents a new method to assess the entire USL and the additional support structures *in situ* rather than *ex vivo*, providing crucial mechanical data about the role the USL plays in USLS suspension. This testing methodology allowed us to test the weakest region of the USL, the cervical region, since the force was applied directly on this region through umbilical tape. During testing, the USL experienced not only tension, but inevitably also compression and shear given the geometry of the ligament and its position relative to the applied load. Moreover, to ensure that the USL was the primary anatomical structure being pulled, we secured the surrounding pelvic tissues to the base plate of the testing machine using a strong adhesive tape. Only the vaginal vault and USL structures were left exposed for testing. Although this prevented significant loading of other anatomical structures, other pelvic tissues were likely loaded and stretched together with the USL. The loads reported here may thus overestimate the loads that are experienced by the USL alone, without their connections to the pelvis. Despite these limitations, the load values we reported fall within the values reported in the literature when loading the rat vagina and USL attachments together *in vivo*^[63]^ and uniaxially testing the isolated rat USL *ex vivo*.^[75]^ The load-displacement curves are very similar to those reported by Donaldson and De Vita, both qualitatively and quantitatively.^[75]^

### 2.6 Study advantages and limitations

The principal strength of this study was the use of a prolapse surgical model to investigate a potential therapeutic for prolapse repair. In the last year, the first study to describe implantation of a mesh biomaterial in the rodent vagina was published,^[66]^ but moving away from mesh-based repairs is advantageous to explore a wide array of possible therapeutics. Other studies investigating materials for prolapse repair utilized abdominal wall hernia models which are poorly predictive of material compatibility with the pelvic organs.^[42,44]^ Next, the *in situ* mechanical testing of the USL-vaginal interface following suture-only repair and repair with augmentation provides measurable outcomes to assess the therapeutic potential of the implanted material for POP. The mechanical testing protocol described here can also be adjusted for use in other pelvic floor models, expanding the range of tools available to researchers to measure outcomes of new biomaterial interventions. Last, the surgical procedures were performed by one surgeon rather than multiple surgeons.

However, this study was subject to several limitations. Controlling for parity of the vendor available animals had the major drawback of receiving animals without exact age information. Without information regarding how recently the rats had delivered their second litters, it was not possible to control for animal weight. Additionally, the rodent pelvis has a horizontal orientation that does not replicate the gravitational effects seen in humans or non-human primates. This and the lack of spontaneous prolapse in the rat models does limit some applicability of these results to humans, but the use of multiparous rats is a strength of this work since this accounts for the leading risk factor in the development of POP.^[76]^ Last, although it is ideal to have a study with a single surgeon, the homogeneity in surgical technique does not capture the variability in outcomes from a diverse population of surgeons.^[77]^

## 3. Conclusions

With this study we describe the first hydrogel augmentation of USLS in rodents, demonstrating the usefulness of the USLS rat model in the investigation of therapeutics for prolapse treatment. With a reoperation rate following prolapse surgery of up to 30%^[15,78,79]^ and major drawbacks to the use of mesh-based biomaterials, there is an urgent need explore alternative biomaterials with properties that are more compatible with the vagina and pelvic floor. The USLS model provides the means for these materials investigations. We were able to assess the *in vivo* efficacy of our supramolecular fibrous hydrogel composite to augment USLS repair in our newly developed animal model. With a carrier hydrogel to encapsulate the guest-host fibers, the material was used to augment the USL-vaginal vault interface created during the USLS procedure. Applying an *in situ* tensile testing method, the repair with hydrogel augmentation significantly increased the pull-out load (∼1.4 N) compared to the USLS procedure with sutures alone (∼ 0.9 N). This demonstrated that the hydrogel assisted in recovery of mechanical properties to be closer to the intact USL tissue (pull-out force ∼ 1.7 N). *In vivo* hydrogel degradation tracking demonstrated exponential decay of fluorescent signal resulting in near dissolution of the hydrogel at 6 weeks post-surgery (∼ 94% degraded), further supporting the premise that superior tissue healing and integration accounts for the recovery of tensile resistance. Together, these results support the potential of this hydrogel platform to augment USLS procedures.

## 4. Experimental Section

### 4.1 Ad-MeHA and CD-MeHA hydrogel synthesis

Hyaluronic acid (HA) was functionalized with photoreactive methacrylates (Me)^[80]^ and then either adamantane (Ad) or β-cyclodextrin (CD)^[49]^ as previously described. Briefly, the HA backbone was methacrylated to produce methacrylate-modified HA (MeHA) via esterification with the primary hydroxyl group of sodium HA at pH 8-9. Next, the MeHA was reacted with proton exchange resin and titrated with tert-butyl ammonium salt (TBA)-OH to yield methacrylated HA-TBA (MeHA-TBA). Ad-modified MeHA (Ad-MeHA) and β-CD-modified MeHA (CD-MeHA) were then synthesized by anhydrous coupling. In Ad-MeHA synthesis, MeHA was modified with 1-adamantane acetic acid via di-tert-butyl bicarbonate (BOC_2_O)/4-dimethylaminopyridine (DMAP) esterification. Separately, CD-MeHA was prepared by coupling 6-(6-aminohexyl)amino-6-deoxy-β-cyclodextrin (CD-HDA) to HA via (benzotriazol-1-yloxy) tris(dimethylamino) phosphonium hexafluorophosphate (BOP) amidation. Synthesis products were dialyzed against deionized water, frozen, and lyophilized. The degree of HA modification with Me was controlled by the amount of methacrylic anhydride introduced during synthesis and was determined to be 16%. Modification of the HA backbone with Ad and CD was determined to be 16.5% (Figure S5) and 16% (Figure S6) respectively, with all percentages calculated by ^1^H NMR.

### 4.2 Solid-phase peptide synthesis

Using a Liberty Blue automated microwave-assisted peptide synthesizer, peptides were synthesized via solid-phase synthesis as described previously.^[81,82]^ Briefly, rink Amide MBHA high-loaded (0.78 mmol/g) resin was used along with solid-supported Fmoc-protected amino acid residues. The resin was swelled with 20% (v/v) piperidine in DMF and the amino acids were sequentially added from C to N-terminus. The resulting peptides were collected and cleaved in 92.5% trifluoroacetic acid, 2.5% triisopropylsilane, 2.5% (2,2-(ethylenedioxy)diethanethiol), and 2.5% water for 2 h and filtered to separate the resin. Two methacrylate-modified peptides were synthesized for use in the hydrogel system: an MMP-degradable peptide (MeP, Methacrylate-GGNS-VPMS↓MRGG-GNCG, Figure S7) and carrier peptide (Methacrylate-GKKCG) for later fluorophore conjugation. A Cy-7.5 fluorophore (cyanine7.5-maleimide, Lumiprobe) was conjugated to the carrier peptide via thiol-maleimide click chemistry, mixing equal molar amounts in 1x PBS for 2 h. The final product was a Cy7.5 labeled methacrylated peptide (Cy7.5-Me). Peptides were precipitated in cold ether, dried overnight, resuspended in water, frozen, and lyophilized. Synthesis was confirmed via matrix-assisted laser desorption/ionization (MALDI).

### 4.3 MePHA hydrogel synthesis

Methacrylated degradable peptide (MeP)-modified HA (MePHA) was synthesized as previously described.^[82]^ Briefly, the HA backbone was modified with maleimide (Ma) groups to facilitate the aqueous addition of the MeP via thiol-maleimide “click” chemistry. In the first step, maleimide HA (MaHA) was synthesized by reacting aminated maleimide salt with tetrabutylammonium-HA (HA-TBA) via a BOP coupling agent to form an amide linkage between the carboxyl group of HA and the amine group of the maleimide salt. The thiolated peptide (containing a cysteine residue) was synthesized with a terminal methacrylate group to allow for later UV light-initiated crosslinking. ^1^H NMR confirmed 22% maleimide modification of the HA backbone (Figure S8) followed by successful conjugation with the MeP to produce MePHA (Figure S9).

### 4.4 Electrospinning

Ad-MeHA and CD-MeHA were dissolved at 2% (w/v) in DI water along with 3.5% (w/v) poly(ethylene oxide) (PEO, 900 kDa) and 0.05% (w/v) Irgacure 2959 (I2959) for 24-48 h prior to electrospinning. The polymer solutions were electrospun using an Elmarco NanoSpider (NS Lab) with the following collection parameters: applied voltage: ∼ 45 kV, electrode distance: 22 cm, orifice diameter: 0.7 mm, substrate: Teflon paper, substrate speed: 20 mm min^-1^, carriage speed: 100 mm sec^-1^. Hydrogel nanofibers were deposited onto Teflon paper, placed into a container which was purged with nitrogen, and crosslinked with UV light (254 nm) for 10 min (VWR Crosslinker Box, 115 V).

### 4.5 Hydrogel formulations

Our lab has previously described the process of collecting and preparing the guest and host electrospun fibers.^[49]^ Briefly, guest Ad-MeHA and host CD-MeHA fibers were hydrated (0.1% w/v in DI water) overnight at 37°C to remove PEO from the electrospinning process, centrifuged, supernatant discarded, and then lyophilized. Once dry, the fibers were again hydrated at 0.1% w/v in DI water, allowed to swell at 37°C for at least 2 h, and then passed through needles of progressively smaller gauge sizes (16G-30G) via trituration. This process separated any adjoined fibers and resulted in a reproducible fiber suspension. MePHA (6 w/v%) was added to envelope the Ad-MeHA/CD-MeHA mixed fibers along with 1 mM lithium acylphosphinate (LAP) to allow for UV light initiated polymerization (365 nm, 10 mW cm^-2^). The final formulation resulted in 70 v/v% of fibers and 30 v/v% of the MePHA with LAP solution.

### 4.6 Rheology

All rheological measurements were performed on an Anton Paar MCR 302 rheometer with the plate temperature set at 37°C as described previously.^[49]^ The hydrogel formulation was tested using a parallel plate (PP08-S; 8 mm diameter, sand blasted) geometry and 25 μm gap distance. Injectability of the hydrogel formulation was tested via a cyclic deformation test alternating between 0.5% and 250% strain to verify shear-thinning and self-healing capabilities. Next, a time sweep (1 Hz, 0.5% strain) with UV crosslinking (365 nm, 10 mW cm^-2^, 2 min) was performed.

### 4.7 Cytocompatibility and hemocompatibility evaluations

Evaluation of cytocompatibility and hemocompatibility was conducted using an Alamar blue assay (Invitrogen) with human mesenchymal stromal cells (hMSCs) and via material incubation with rat red blood cells (RBCs), respectively. *In vitro* testing of the hydrogel cytotoxicity followed ISO 10993 standards where the hydrogel components were added to cell culture media (1% w/v) and then applied to a monolayer culture of hMSCs. Cells were seeded at 5000 hMSCs per well in a 48-well plate with the Alamar blue applied at time intervals of 1 and 3 days of culture. Metabolically active cells reduce the Alamar blue reagent to a fluorescent byproduct (resorufin). Fluorescent signal was read by a plate reader at 565 nm (excitation), 595 nm (emission) after incubation for 5 h in the dark. Hemolysis testing followed ISO 10993 and ASTM F756 standards where blood was harvested immediately prior to assay preparation. Fresh blood was suspended in 0.01% heparin (Sigma Aldrich) in PBS. The mixture was immediately centrifuged at 1000 rpm for 15 min and then washed with PBS until the supernatant above the pellet of RBCs was clear. The RBCs were then diluted using PBS and incubated for 1 h at 37°C with the hydrogel components. After incubation, the RBCs were collected and centrifuged again at 1000 rpm for 15 min before the absorbance of the supernatants were read at 540 nm using a plate reader. The results were calculated as hemolysis (%)

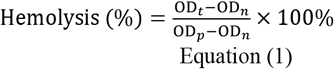

where OD_t_, OD_n_, and OD_p_ were the absorbance values of the samples, negative control (PBS), and positive control (0.1% Triton X-100), respectively.

### 4.8 Animal care

This study was conducted in compliance with the Animal Welfare Act, the Implementing Animal Welfare Regulations, and in accordance with the principles of the Guide for the Care and Use of Laboratory Animals. The University of Virginia Animal Care and Use Committee approved all animal procedures. A total of 24 multiparous (2 litter) female Lewis rat breeders (Charles River Laboratories) weighing 274.8 ± 19.3 g were pair housed in a vivarium accredited by the American Association for the Accreditation of Laboratory Animal Care and provided with food and water *ad libitum*. Animals were between 4 and 6 months of age to accommodate the 2 litter requirement. Animals were maintained on a 12-hour light-dark cycle.

### 4.9 Anesthesia, pain management, and antibiotics

All animal-related details were described in detail previously.^[32]^ Briefly, animals were anesthetized via isoflurane and the surgical site was aseptically prepared by repeated washes with alcohol and iodine. The depth of anesthesia was monitored by the response of the animal to a slight toe pinch, where the lack of response was considered the surgical plane of anesthesia. Core temperature was maintained using a heated water perfusion system. Rats were administered slow-release buprenorphine (Bup XR – 72 h; 1.3 mg kg^-1^, subcutaneously), slow-release meloxicam (72 h; 1.0 mg kg^-1^, subcutaneously), and baytril (10 mg kg^-1^, subcutaneously) prior to surgery. Since meloxicam is known to cause dehydration, rats were also administered 0.9% sterile saline (10 mL kg^-1^, subcutaneously). Animal pain and distress were monitored daily by qualified members of the veterinary staff to determine the need for additional analgesia.

### 4.10 Surgical procedures

Using aseptic technique, a vertical midline skin incision was made along the abdomen and the fascia was separated to expose the underlying abdominal cavity. Hysterectomy was performed, where the uterine horns were trimmed after separation from the ovaries and fat pads, leaving the cervix, vagina, and its support tissues intact. One delayed absorbable suture (3.0 polydiaxanone, PDS II, Ethicon) was placed through the vaginal vault and the exposed uterosacral ligament (USL) bilaterally. Tying the sutures down simultaneously closed the vaginal vault and elevated it toward the USLs. A single uterosacral stitch was used due to limited space along the USL in the rat model and our goal to have spatial ability to perform mechanical testing of the ligament. For hydrogel augmentation, material components were sterilized separately and then prepared aseptically. A total of 40 μL (20 μL each side) hydrogel precursor was delivered over the sutures and then UV crosslinked (365 nm, 10 mW cm^-2^, 2 min). Animals in the sham experimental group were used for healthy animal controls to assess normal tissue properties. Animals were randomly assigned to one of the following experimental groups: sham surgery with no prolapse repair (USL; *n* = 8), prolapse repair (USLS; *n* = 8), or prolapse hydrogel repair (USLS+H; *n* = 8).

### 4.11 *In vivo* hydrogel degradation analysis

To evaluate *in vivo* hydrogel degradation, hydrogel nanofibers were labeled with the Cy7.5-Me peptide via UV light-initiated conjugation in the presence of LAP photoinitiator. The fluorophore was covalently bound via the methacrylates present on the surface of the hydrogel fibers during photocuring. Serial images were taken during a 6 week period postoperative using a LagoX live imaging system (excitation: 770 nm, emission: 810 nm) with signal intensity measured by integrating equivalent areas over the region of interest. Quantified signal was normalized to peak intensity for individual hydrogel boluses immediately following surgery and then averaged to obtain degradation profiles for each hydrogel (*n* = 4).

### 4.12 Mechanical testing of the USLS and prolapse repair junction

At 24 weeks post-surgery, the uterosacral ligament (USL) or USLS prolapse repair junction (USLS, USLS+H) were prepared for mechanical testing using a single column Instron (5943 S3873) universal testing system. A 10 N load cell was used for testing. Methods were described, including a video demonstration, in previous work.^[32]^ Briefly, umbilical tape (Ethicon U10T) was threaded beneath the suture/new tissue formation between the vaginal vault and the uterosacral ligament. The tissue was pre-loaded at 0.015 N and then preconditioned at an elongation rate of 0.1 mm/s for 1 min. The tissue was then pulled to failure at the same elongation rate.

### 4.13 Statistical analysis

All data are presented as means and their standard deviations (SDs). The data were statistically analyzed by one-way ANOVA tests with multiple comparisons using GraphPad Prism 9.0. *P* values < 0.05 were considered statistically significant where \**P* < 0.05, \*\**P* < 0.01, \*\*\**P* < 0.001 and \*\*\*\**P* < 0.0001.

## Supporting Information

MALDI and ^1^H NMR spectra for the hydrogel components as well as additional data from the mechanical testing experiments can be found in the Supporting Information.

## Acknowledgments

We thank Luna Innovations for the use of their NanoSpider for hydrogel nanofiber production, Prof. Rachel Letteri for the use of her peptide synthesizer, and Prof. George Christ for use of his surgical space. This work was supported by the UVA-Coulter Translational Research Partnership and the DoD (W81XWH-19-1-0157).

