## Supporting Information for "Supramolecular fibrous hydrogel augmentation of uterosacral ligament suspension for treatment of pelvic organ prolapse"

W. Wolfe
Scripps Institution of Oceanography
University of California, San Diego
La Jolla, CA 92093, USA

J.L. Gentry, C.B. Highley, S.R. Caliari
Department of Biomedical Engineering
University of Virginia
Charlottesville, VA 22903, USA

R. De Vita
Stretch Lab
Department of Biomedical Engineering and Mechanics
Virginia Tech
Blacksburg, VA 24061, USA

M.H. Vaughan
Department of Obstetrics and Gynecology
University of Virginia
Charlottesville, VA 22903, USA


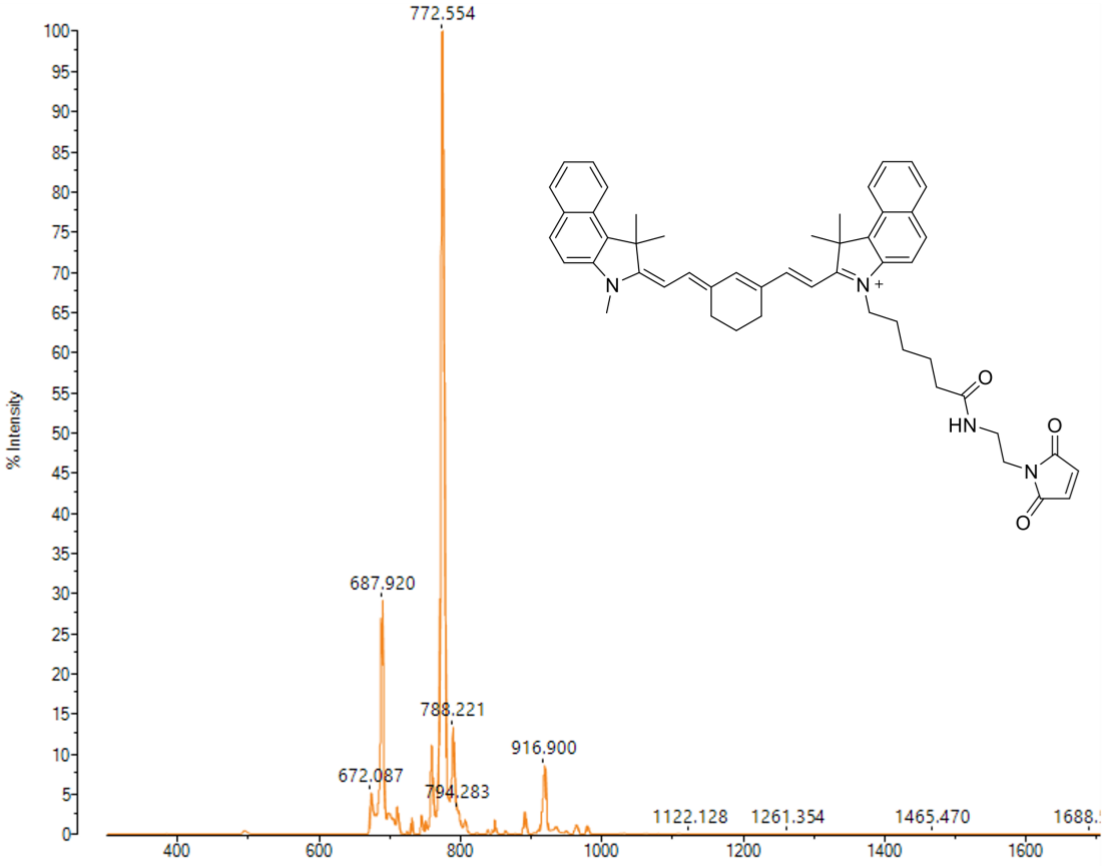


**Figure S1: MALDI spectrum of maleimide-modified cyanine 7.5 (Cy7.5) fluorophore.** Purchased from Lumiprobe Corporation. Expected mass: 771.4 g/mol. Actual mass: 772.5 g/mol.

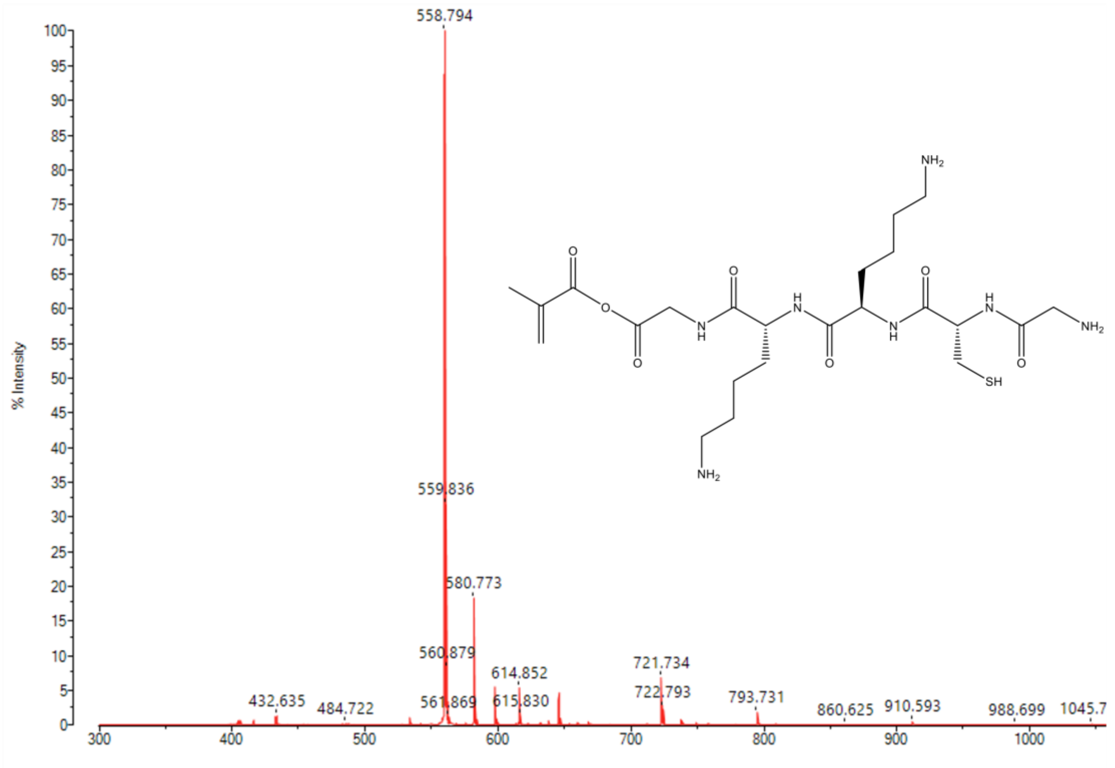


**Figure S2: MALDI spectrum of methacrylated carrier peptide.** Methacrylated peptide with sequence methacrylate-GKKCG synthesized for conjugation (via the thiol on the cysteine residue) with maleimide-modified Cy7.5 fluorophore. Expected mass: 559.2 g/mol. Actual mass: 558.8 g/mol.

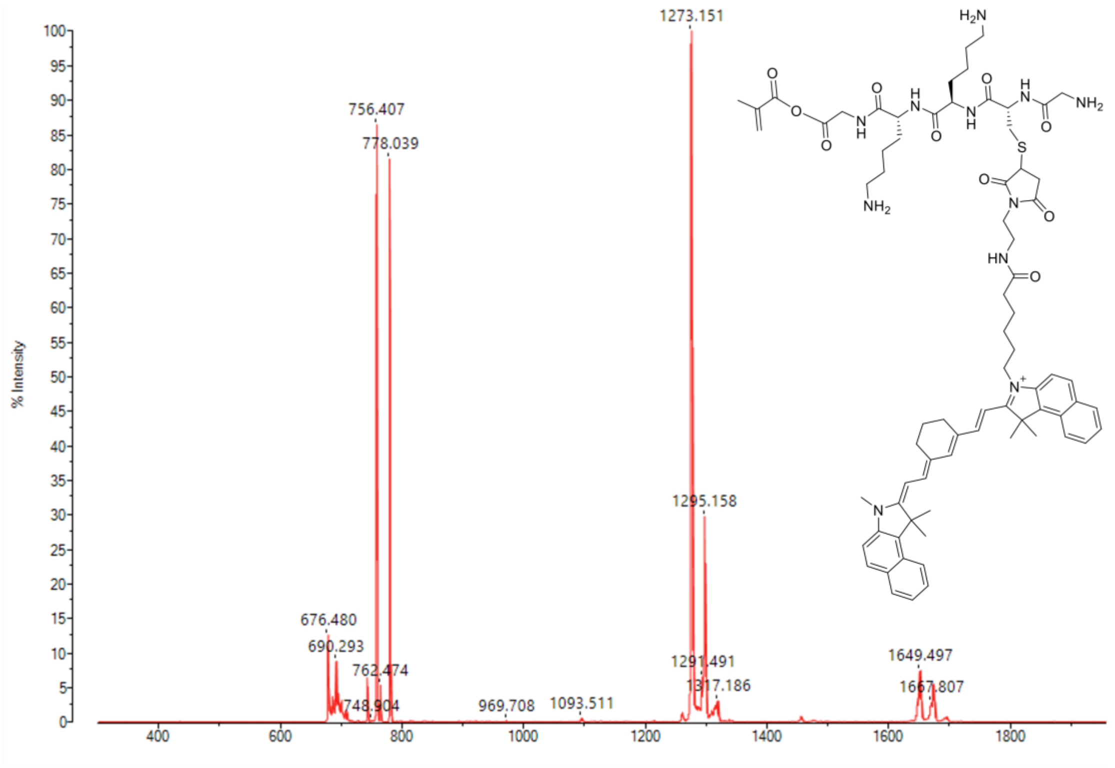


**Figure S3: MALDI spectrum of the Cy7.5-labeled methacrylated peptide.** Methacrylated peptide conjugated with maleimide-modified Cy7.5 fluorophore to enable covalent conjugation to the HA backbone via photopolymerization of methacrylates. Secondary peaks show unreacted fluorophore. Expected mass: 1273.6 g/mol. Actual mass: 1273.1 g/mol.

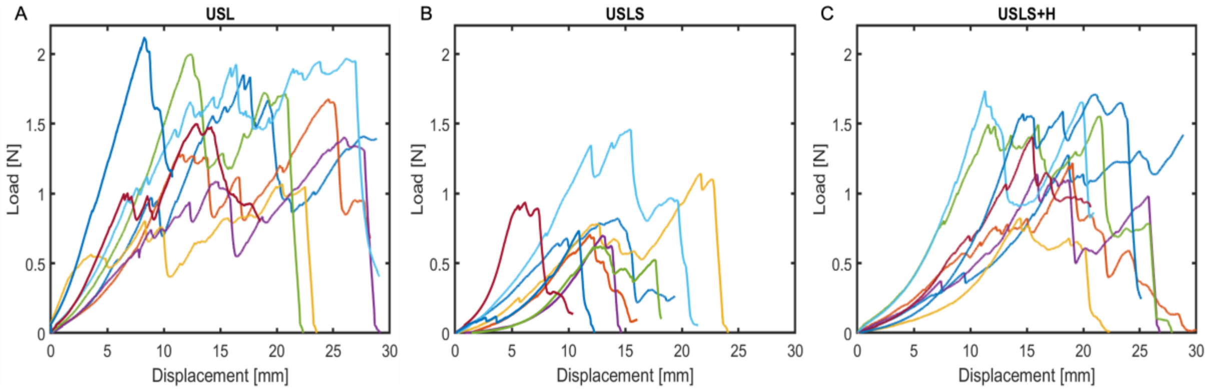


**Figure S4: Load-displacement curves from mechanical testing to failure.** Data are separated by experimental groups A) intact USL, B) USLS, and C) USLS with the hydrogel augmentation (USLS+H). $n$ = 8 animals per experimental group.

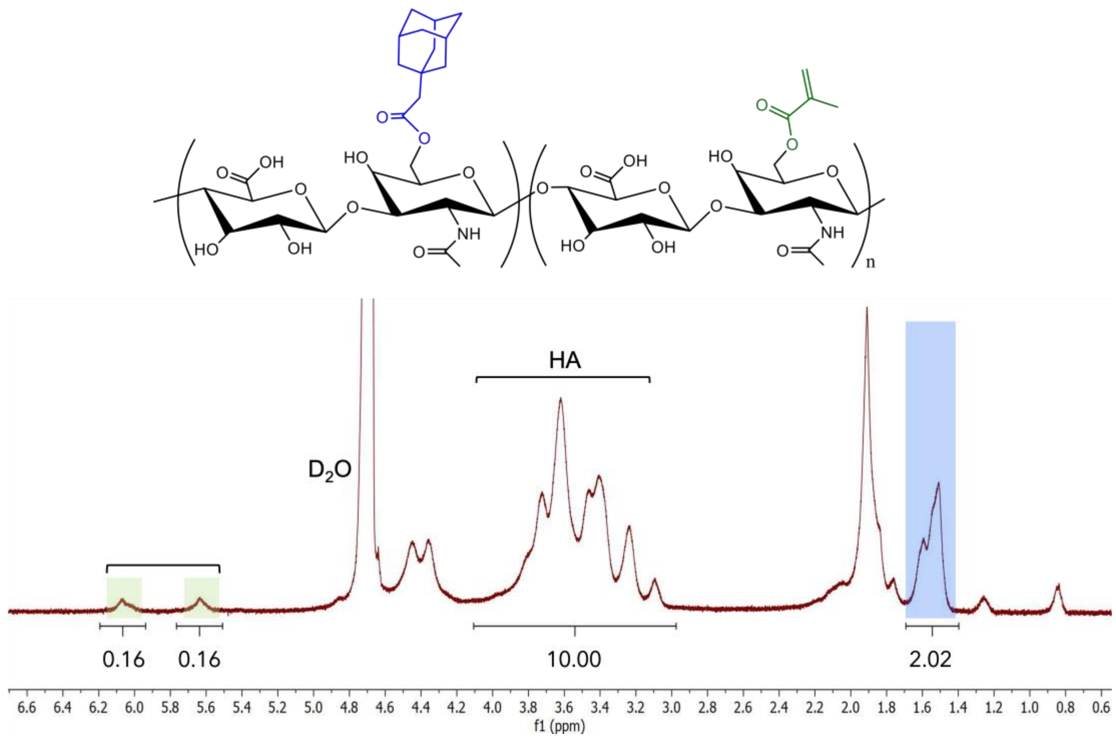


**Figure S5**: **$^1$H NMR spectrum of adamantane and methacrylate-modified hyaluronic acid (Ad-MeHA).** The degree of modification, relative to the HA backbone (δ = 3.10- 4.10, 10 H), was determined to be 16.5% for the adamantane and 16% for the methacrylate. Adamantane modification was determined via integration of the ethyl multiplet of adamantane, δ = 1.40 – 1.70, 12 H (highlighted blue) while methacrylate functionalization was determined via integration of the vinyl group in the methacrylate, δ = 5.82, 1 H and δ = 6.25, 1 H (highlighted green).

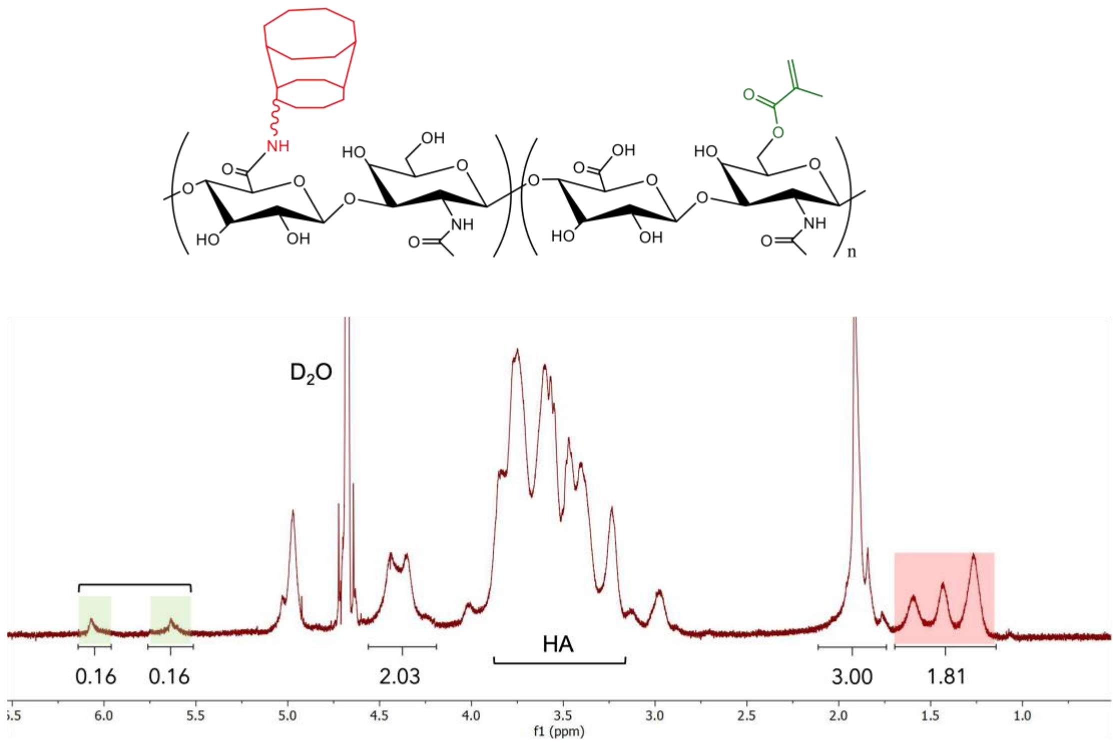


**Figure S6**: **[1]H NMR spectrum of β-cyclodextrin and methacrylate-modified hyaluronic acid (CD-MeHA).** The degree of modification, relative to the methyl singlet of HA (δ = 2.1, 3 H), was determined to be 16% for the β-cyclodextrin and 16% for the methacrylate. β-cyclodextrin modification was determined via integration of the hexane linker, δ = 1.20-1.75, 12 H (highlighted red) while methacrylate functionalization was determined via integration of the vinyl group in the methacrylate, δ = 5.82, 1 H and δ = 6.25, 1 H (highlighted green).

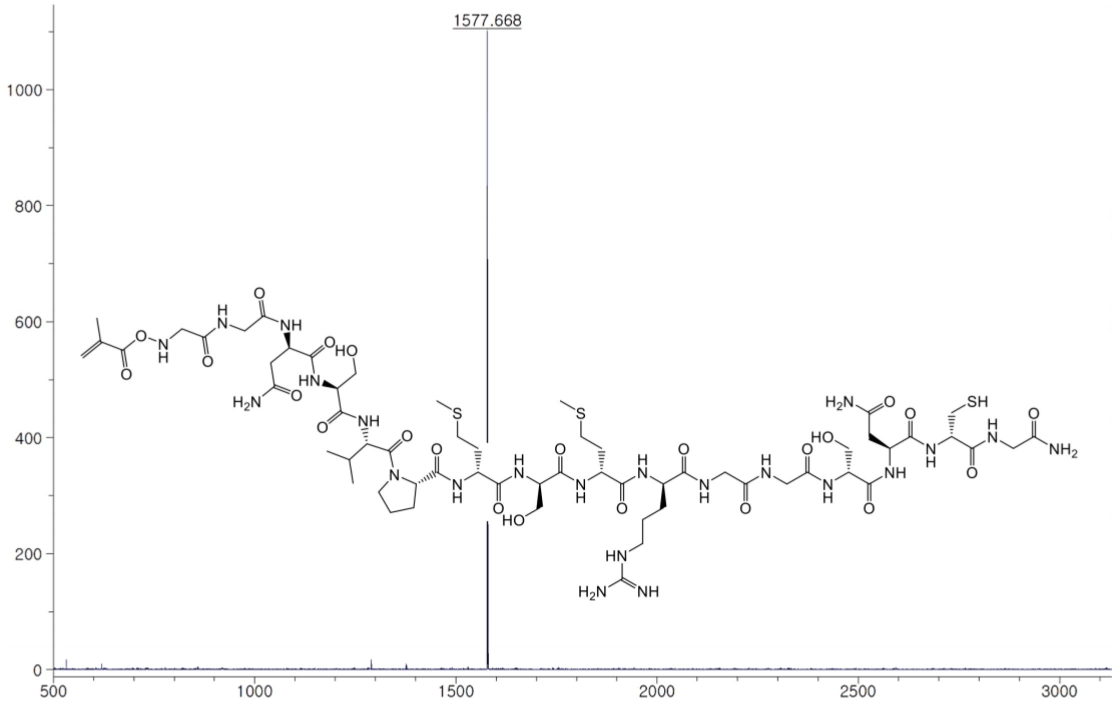


**Figure S7: MALDI spectrum of methacrylated MMP-degradable peptide (MeP).** Degradable peptide with sequence methacrylate-GGNS-VPMS↓MRGG-GNCG. Expected mass: 1578 g/mol. Actual mass: 1577.6 g/mol.

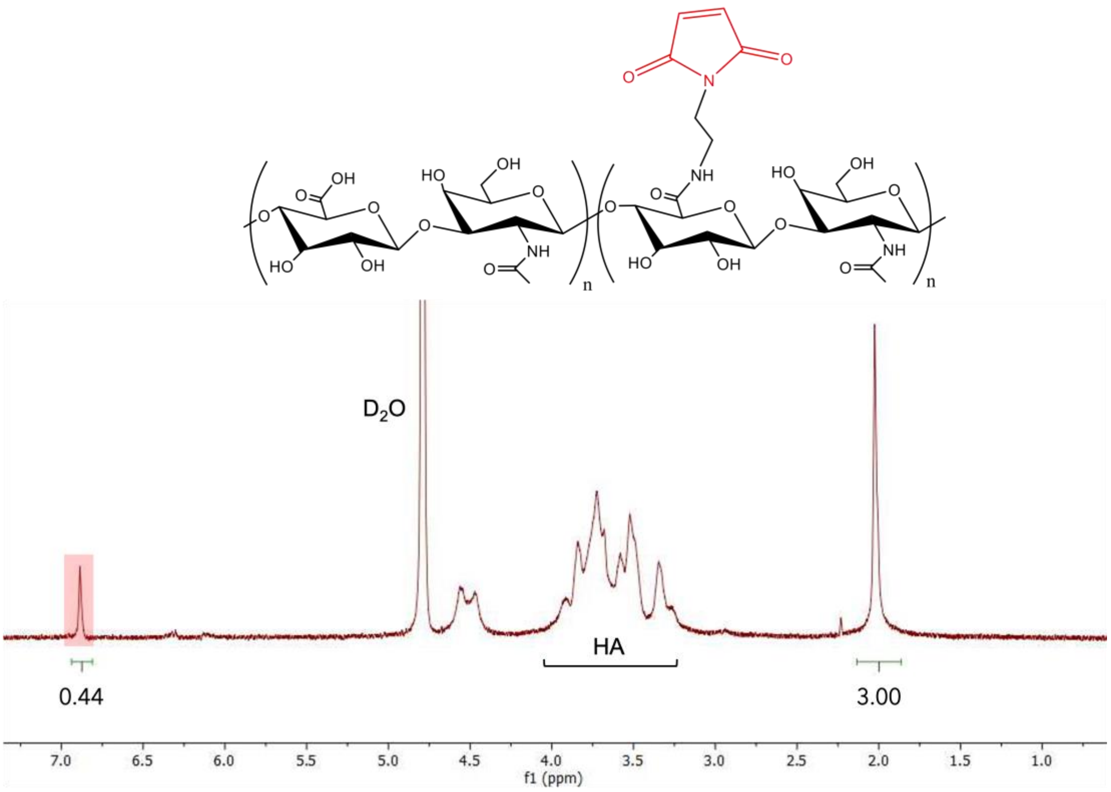


**Figure S8: $^1$H NMR spectrum of maleimide-modified hyaluronic acid (MaHA)**. Maleimide functionalization was determined to be 22% from integration of the doublet δ = 6.92, 2 H (highlighted red) relative to the methyl singlet of HA (δ = 2.1, 3 H).

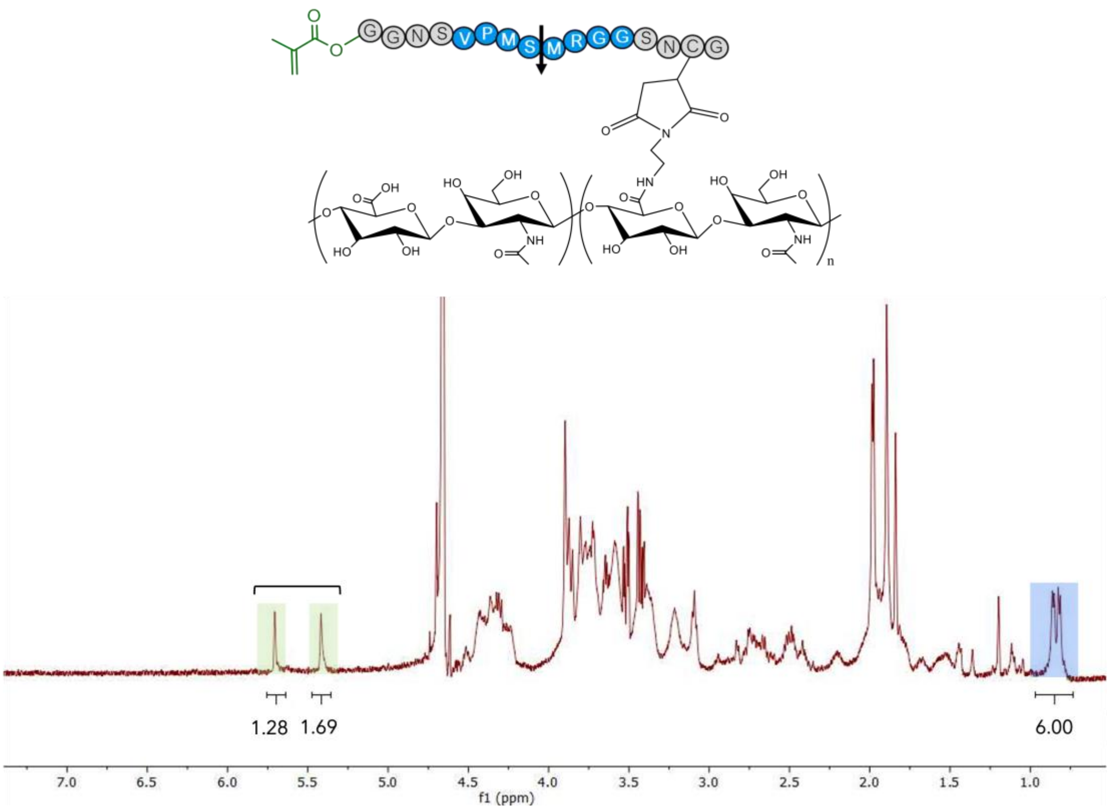


**Figure S9: [1]H NMR spectrum of methacrylated peptide-modified hyaluronic acid (MePHA)**. The MMP-degradable thiolated peptide (MeP) was conjugated to the maleimide-modified HA (MaHA) to form methacrylate peptide-modified hyaluronic acid (MePHA) that was both photocrosslinkable and MMP-degradable. Loss of the maleimide peak, and the corresponding introduction of methacrylate peaks, indicated successful peptide conjugation. Assuming initial maleimide functionalization of 22% (from Fig. S8), integration of the methacrylate peaks ($\delta$ = 5.82, 1 H and $\delta$ = 6.25, 1 H; highlighted green) relative to the valine peak of the peptide sequence ($\delta$ = 1.02, 6 H; highlighted blue), in combination with disappearance of maleimide peaks, indicates successful and efficient conjugation of MeP to MaHA.
